# BRD4 inhibitors block telomere elongation

**DOI:** 10.1101/112169

**Authors:** Steven Wang, Alexandra M. Pike, Stella S. Lee, Carla J. Connelly, Carol W. Greider

**Author notes:** Present Address: Stella S. Lee, Office of the Commissioner, Food and Drug Administration, Silver Spring, MD, 20993, USA.

## Abstract

Cancer cells maintain telomere length equilibrium to avoid senescence and apoptosis induced by short telomeres, which are triggered by the DNA damage response. Limiting the potential for telomere maintenance in cancer cells has been long been proposed as a therapeutic target. Using an unbiased shRNA screen targeting known kinases, we identified bromodomain 4 (BRD4) as a telomere length regulator. Four independent BRD4 inhibitors blocked telomere elongation, in a dose dependent manner, in mouse cells overexpressing telomerase. Long-term treatment with BRD4 inhibitors caused telomere shortening in both mouse and human cells, suggesting BRD4 plays a role in telomere maintenance *in vivo*. Telomerase enzymatic activity was not directly affected by BRD4 inhibition. BRD4 is in clinical trials for a number of cancers, but its effects on telomere maintenance have not been previously reported investigated.

## INTRODUCTION

Telomere length maintenance is required for long-term division of cells. When telomeres become short, they no longer protect chromosome ends, and cells undergo senescence or apoptosis (1-3). This requirement for telomere length maintenance suggested that blocking telomere elongation might block the growth of cancer cells in some settings (4). Telomerase elongates telomeres and maintains a telomere length equilibrium that prevents telomeres from becoming critically short (5). In the setting of insufficient telomerase or other telomere gene mutations, human short telomeres syndromes manifest as bone marrow failure, immunodeficiency, enteropathy, pulmonary fibrosis and emphysema (6,7). Conversely, most cancer cells upregulate telomerase (8-10). Recent experiments have shown promoter mutations that increase the expression of the telomerase catalytic component TERT, are very common in many cancers (11,12). Indeed, germline mutations in the TERT promoter, or POT1, predispose to familial melanoma, glioma or CLL respectively (13-15). This suggests there may be a long telomere state that is cancer prone (7).

Telomerase inhibitors have been proposed as potential cancer therapeutics for over 25 years (4,10,16,17). BIBR1532 is a potent inhibitor of telomerase in cell extracts and in cell culture (18), but has limited solubility (19,20) and has not progressed to clinical trials. Imtelstat is an anti-sense molecule that inhibits telomerase by binding to the intrinsic RNA template. Imtelstat is an effective *in vitro* telomerase inhibitor and it shortens telomeres in human cultured cells (21,22). However, it has failed phase II clinical trials and the mode of action in some malignancies may be due to off-target effects (21,23,24). With our current understanding of short and long telomere syndromes, and a growing understanding of cancers that rely on telomerase, it is possible to revisit the concept of telomere shortening in cancer with a more nuanced, approach.

Several groups identified telomerase regulators through direct screening of compounds or genes that block telomerase enzyme activity (25-27). We took a different approach by identifying pathways that might block telomere elongation without direct inhibition of telomerase activity or telomerase transription. Telomere elongation is regulated by shelterin proteins (28-30) and by post-translational modification (31-35). To identify kinase pathways that might regulate telomere length, we designed an unbiased shRNA screen against kinases.

In this screen, we identified BRD4 as a novel, positive regulator of telomere length. BRD4 is a BET family protein that contains a bromodomain, which binds to acetylated lysines (36). It also has histone acetyl transferase activity (37), and kinase activity (38). BRD4 is a pleiotropic protein with roles in cell cycle regulation, chromatin structure, and transcriptional regulation (39). BRD4 has not previously been implicated in telomere length regulation. Because several BRD4 inhibitors are currently in clinical trials for cancer, understanding their potential effects on telomere length will be important to discover both its mechanism of action and potential side effects.

## MATERIAL AND METHODS

### Cell Culture

HeLa and mouse fibroblasts were cultured in DMEM (Gibco) with 1% penicillin/streptomycin/glutamine (PSG) and 10% heat-inactivated fetal bovine serum (Gibco). Drugs were dissolved in DMSO and added to cell culture media at indicated concentrations.

### Lentiviral shRNA kinase library

Decode Pooled Human GIPZ Kinase Library (GE Dharmacon RHS6078) was used to screen for telomere length regulators. This library contained 4675 shRNAs, in pGIPZ lentiviral vectors, directed against 706 kinase and kinase related genes. HeLa cells were transduced with 500-fold representation of the library at an MOI of 0.1. Experiments were performed in triplicate. After transduction, cells were cultured cells for 7 weeks to allow changes in telomere length over many cell divisions.

### Telomere Flow-FISH and fluorescence activated cell sorting

We adapted a version of telomere Flow-FISH (40) to sort cells with short telomeres. 4×10^7^ cells were fixed in 1.5% paraformaldeyhyde for 10 minutes then dehydrated in 100% methanol overnight. Cells were washed with PBS, hybridized with probe and washed, as described (40). Cells were resuspended at a concentration of 5×10^6^/mL in modified propidium iodide staining solution, consisting of PBS with 0.1% Triton X-100, 200 μg/mL RNase A (Sigma), 20 μg/mL propidium iodide (Sigma), and 0.1% sodium dodecyl sulfate (SDS, Bio-Rad). SDS was found to inhibit aggregation of fixed cells during cell sorting. Cells were incubated for 30 minutes at room temperature, protected from light. Sorting was performed on a MoFlo cell sorter (Becton Dickinson). Cells were first gated on the G1 cell cycle peak to ensure only cells with 2N ploidy were measured. Telomere FITC signal was measured in the linear range, and the cells with the 7% shortest telomeres were collected. A minimum of 2.5 million cells was collected for each fraction. Unsorted cells were collected as a control.

### DNA preparation, amplification and sequencing

Genomic DNA was harvested from sorted cells by phenol-chloroform extraction (41). shRNA insert was amplified from common flanking regions by PCR as described (42). PCR primers contained indices for multiplexing (42). PCR products were purified with Agencourt AMPure XP Beads (Beckman Coulter), and then sequenced with single end 50 base pair reads using Illumina HiSeq 2000.

### Bioinformatic analysis

Illumina reads were aligned to reference sequences using Bowtie2 as described (42,43), and enriched genes were determined by evaluating 3 biological replicates with MAGeCK analysis (44). MAGeCK first normalizes samples based on median number of reads per sample. Next, it models a relationship for mean number of reads versus variance, based on shRNA read counts across all replicates. Finally, MAGeCK ranks genes by taking into account the enrichment and p-value of each particular shRNA. Since there are multiple shRNAs per gene, genes with multiple highly ranked shRNAs are ranked higher than those with just a few. Statistically significant enriched genes were inspected manually, and prioritized for further characterization based on MAGeCK rank, availability of chemical inhibitors, and known role in potential telomere pathways such as DNA damage, cell cycle regulation, checkpoint regulation, DNA replication and chromatin modification.

### Small molecular inhibitors

KU-55933 (R&D Systems 3544) was used to inhibit ATM. JQ1 (Selleckchem S7110), OTX015 (Selleckchem S7360), I-BET151 (Selleckchem S2780), and MS436 (Selleckchem S7305) were used to inhibit BRD4. Additional inhibitors tested that did not show an effect, are listed in Supplemental table 1.

### Secondary screen for inhibitors of telomere elongation

Immortalized CAST/EiJ fibroblasts were treated with drug or vehicle for 24 hours, then transduced with the SVA lentivirus at MOI=0.5. SVA is a lentiviral vector encoding mTERT and mTR, the catalytic protein and RNA components of telomerase (45). Cells were cultured in the presence of drug or DMSO for 6 days and genomic DNA was isolated at days 2 and 6 post transduction for Southern analysis of telomere length as described (46).

Genomic DNA was digested with MseI (NEB), separated on a 0.7% agarose in 1×TAE (40mM Tris, 20mM Acetate, 1mM EDTA, pH 8.6), denatured in 0.5 M NaOH/1.5 M NaCl and neutralized in 1.5 M NaCl/0.5 M Tris-HCL pH 7.4. DNA was transferred in 20X SSC (3M NaCl, 0.34M NaCitrate) to a nylon membrane (Amersham Hybond N+), crosslinked by UV (Stratagene), and prehybridized for 2 hours in Church buffer (0.5M Sodium Phosphate, pH7.2, 7% SDS, 1% BSA, 1mM EDTA) and hybridized overnight at 65°C with radiolabeled telomere fragment, generated from JHU821 as described, (47) and radiolabeled 2-log DNA ladder (NEB). After washing, the nylon membranes were exposed to Storage Phosphor Screens (GE Healthcare) and scanned on a Storm 825 imager (GE Healthcare). The images were converted using Adobe Photoshop CS6 and adjusted for contrast using the *“*curves” feature within the software. The 2-log ladder marker shown on the left of the Southern blots, indicates the sizes in kb.

### Direct telomerase activity assay

Direct telomerase assay was performed on cell lysates as described with modifications (45). For stable hTR overexpression, pBluescript II SK(+)U1-hTR (48) was cloned into a FUGW-derived lentiviral backbone where Puro was cloned into the GFP site with BamH1 and EcoR1. 293TRE×/FRT cells were transduced with the FUPW-hTR lentivirus and selected for clonal hTR overexpressing cells. Polycistronic TERT, POT1 and TPP1 was flipped into a single genomic FRT site using the Flp-in system (Invitrogen). These cells were treated with DMSO, 5 μM IBET151, 0.5 μM JQ1,25 μ M MS436, or 2.5 μM OT×015 for 48 hours before lysis in 1× CHAPS buffer. Protein was quantified by Bradford assay (Bio-Rad). Equal amounts of protein were incubated with a5 primer (49) and 0.5 mM dTTP, 0.5 mM dATP, 2.92 μM dGTP, and 0.33 μM α32P-dGTP (Perkin Elmer) in telomerase buffer (50 mM Tris-Cl, 30 mM KCl, 1 mM MgCl2, 1 mM spermidine). Reactions were incubated for 15 minutes at 30°C, terminated with stop buffer (20mM EDTA, 10mM Tris), spiked with an end-labeled 18-mer purification control, and telomere products were isolated by phenol-chloroform extraction and ethanol precipitation. Reaction products were separated on a 10% polyacrylamide/7M urea sequencing gel, dried, and imaged on a Storm 825 imager (GE Healthcare). Telomere repeats were quantified in ImageQuant, and processivity values were calculated as described (50). Telomerase catalytic activity was calculated by measuring the intensity of the first telomere repeat normalized to the loading control.

## RESULTS AND DISCUSSION

### BRD4 identified in screen for telomere length regulators

To identify new pathways that regulate telomere length, we carried out an unbiased screen of shRNAs that target kinases in HeLa cells. Kinases are attractive therapeutic targets (51), and there is evidence that they affect telomere length. Loss of Tel1 kinase in yeast, or ATM, its homologue in mammals, results in telomere shortening (33,45,52). CDK1 activity is also required for telomere elongation in yeast (31,32). We screened a lentiviral library of shRNAs, targeting 706 kinase and kinase related genes (GE Dharmacon RHS6078) (42). HeLa cells were transduced with the library in triplicate and cultured for 7 weeks to allow for telomere length changes after multiple divisions. We adapted a Flow-FISH protocol (40) to hybridize telomere probe to telomeres, in intact cells, and then separate cells based on telomere length, using fluorescence activated cell sorting (FACS). Cells with the shortest telomeres were collected and shRNA inserts were amplified, sequenced and genes enriched in the short population, compared to the unsorted population, were identified by aligning to reference sequences with Bowtie2 (43), and ranking gene enrichments with MAGeCK (44). We manually inspected candidates, and prioritized those that were involved in nuclear localization, cell cycle regulation, checkpoints, DNA replication, and chromatin modification. Selected candidates, for which small molecule inhibitors were available, were tested in a secondary screen for inhibition of telomere elongation.

We previously found that inhibition of ATM kinase with the specific inhibitor KU55933, blocked telomere elongation when telomerase was overexpressed (45). Mouse fibroblasts, from a CAST/EiJ strain with short telomeres, transduced with a lentivirus, SVA, encoding both mTR and mTERT, showed robust telomere elongation after 6 days. The ATM inhibitor KU55933 blocked this elongation (45) (Figure 1A). We used this blockage of telomere elongation as a secondary screen to evaluate selected hits from the flow-FISH screen. We tested 36 inhibitors to candidates genes from our screen for their ability to block telomere elongation (Supplementary Table 1). We found that the BRD4 inhibitor, JQ1 (53), effectively blocked telomere elongation (Figure 1A), in a dose dependent manner from 3.3nM to 33 nM (Figure 1B). The degree of inhibition was similar to that seen with the inhibition of ATM by KU55933 (45).

**Figure 1.**
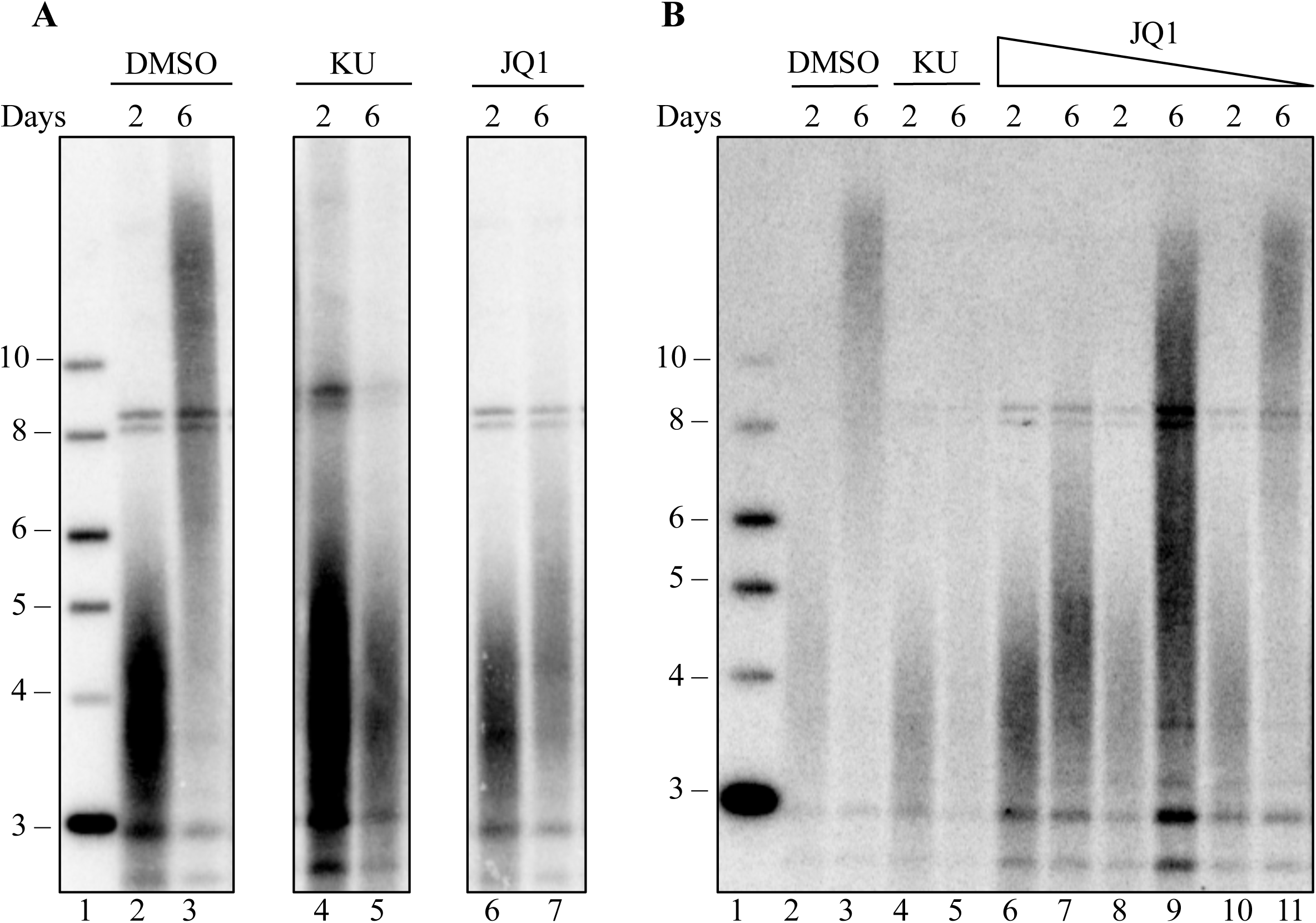
JQ1 blocks telomere elongation in a dose dependent manner. Mouse fibroblasts were transduced with SVA lentivirus, encoding for mTERT and mTR and cultured for six days. (A) Southern blot of mouse fibroblast telomeric DNA, at days 2 and 6 post SVA transduction, grown in the presence of DMSO (lane 2-3), 10 μM KU-55933 (lane 4-5) or 0.1 μ M JQ1 (lane 6-7). Lane 1 shows the 2-log ladder marker (NEB) sizes marked are in in kilobases. (B) Southern blot of mouse fibroblast telomeric DNA, at days 2 and 6 post SVA transduction, in the presence of DMSO (lane 2-3), 10 μ M KU-55933 (lane 4-5), or decreasing concentrations of 33 nM JQ1 (lane 6-7), 11 nM JQ1 (lane 8-9) or 3.3 nM JQ1 (lane 10-11). Lane 1 shows the 2-log ladder marker (NEB) sizes marked are in in kilobases.

We next tested the effects of three additional BRD4 inhibitors, OTX015, I-BET151 and MS436 on telomere elongation. All of these inhibitors blocked telomere elongation, induced by telomerase overexpression, in a dose dependent manner (Figure 2A-C), indicating inhibition is specific to BRD4. All four of these inhibitors, target the bromodomain of BRD4 and interfere with its binding to acetyl lysine thus blocking chromatin binding of BRD4 (54). The effect on telomere elongation is not likely due to transcriptional regulation of telomerase components directly since, in this screen, mTERT and mTR are highly overexpressed from exogenous promoters.

**Figure 2.**
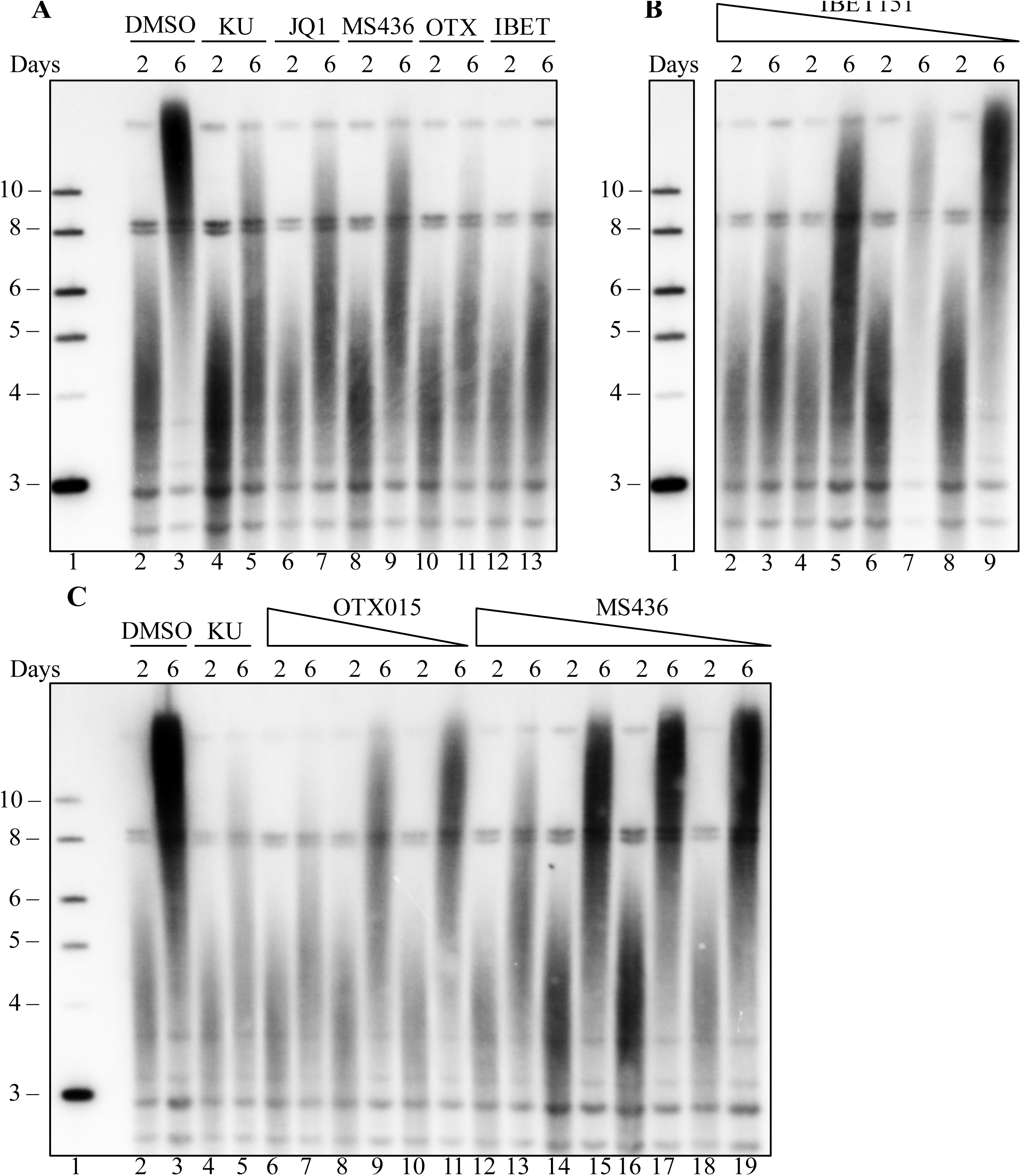
Three additional BRD4 inhibitors block telomere inhibition in a dose dependent manner. Mouse fibroblasts transduced with SVA lentivirus were treated with four different BRD4 inhibitors: JQ1, IBET151, MS436 OTX015. (A) Southern blot of mouse fibroblast telomeric DNA, at days 2 and 6 post SVA transduction, grown in the presence of DMSO (lane 2-3), 10 μ M KU-55933 (lane 4-5), 0.1 μ M JQ1 (lane 6-7), 5 μ M MS436 (lane 8-9), 0.5 μ M OTX015 (lane 10-11), or 1 μM IBET151 (lane 12-13). (B) Dose dependence of IBET151. Southern blot of mouse fibroblast telomeric DNA, at days 2 and 6 post SVA transduction, in the presence of 1 μM (lane 2-3), 0.5 μM (lane 4-5), 0.25 (lane 6-7), or 0.125 μ M (lane 8-9) IBET151. (C) Dose dependence of OTX015 and MS436. Southern blot of mouse fibroblast telomeric DNA, at days 2 and 6 post SVA transduction, in the presence of DMSO (lane 2-3), 10 μM KU-55933(lane 4-5), 250 nM (lane 6-7), 125 nM (lane 8-9) or 62.5 nM OTX015 (lane 10-11), 5 μM (lane 12-13), 2.5 μ M (lane 14-15), 1.25 μ M (lane 16-17) or 0.625 μ M (lane 18-19) MS436. Lane 1 in all panels shows the 2-log ladder marker (NEB), sizes marked are in in kilobases. In Panel B, other lanes between the marker and IBET treated lanes were removed.

### BRD4 does not affect telomerase enzymatic activity

To examine whether the BRD4 inhibitors had a direct effect on telomerase enzyme activity, we used a quantitative direct activity assay (45). 293T cells overexpressing telomerase components were treated with the highest tolerated dose of each of the four BRD4 inhibitors JQ1, OTX015, I-BET151 and MS436. There was no effect of any of these compounds on telomerase activity or processivity *in vitro* (Figure 3A, B), suggesting that BRD4 inhibitors do not block telomere elongation through direct inhibition of telomerase enzymatic activity.

**Figure 3.**
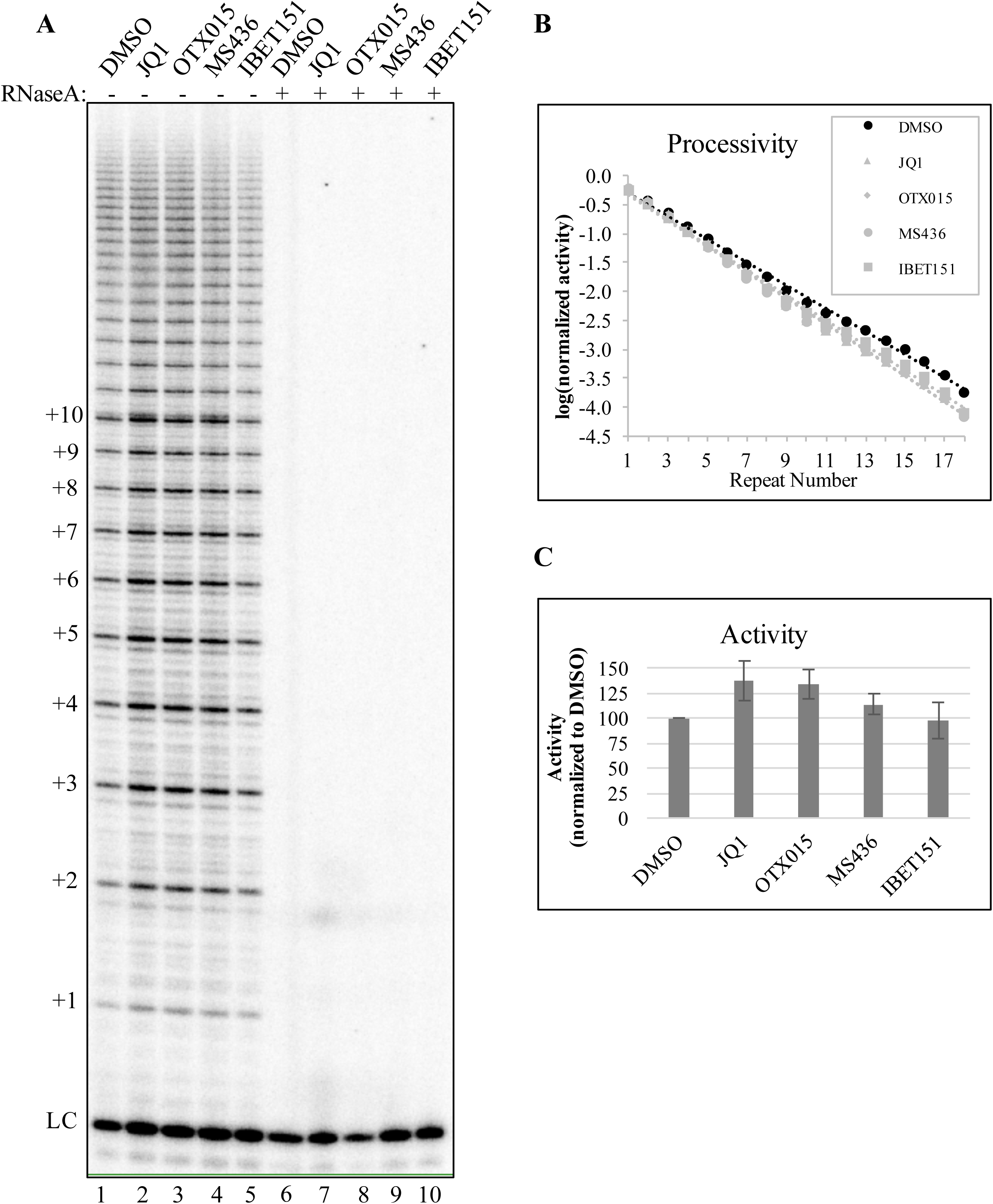
BRD4 inhibition does not affect telomerase enzyme activity. (A) Direct telomerase assay using whole cell lysates of 293TRex cells overexpressing hTR, TERT, POT1, and TPP1, which were treated with DMSO (lane 1), 0.5 μ M JQ1 (lane 2), 2.5 μ M OTX015 (lane 3), 25 μ M MS436 (lane 4), or 5 μ M IBET151 (lane 5). Lanes 6-10 show extracts pretreated with RNase A to show activity due to RNase sensitive telomerase enzyme. (B) Quantification of telomerase processivity and (C) Quantification of telomerase catalytic activity. Values in (B) and (C) are averages of two technical replicates. Error bars in (C) represent the standard deviation.

BRD4 inhibition has pleiotropic phenotypes on the cell cycle by affecting its role as a mitotic bookmark (55), a transcriptional scaffold (55,56) or altering its HAT activity (37). The mechanism by which BRD4 inhibition blocks telomere elongation is not yet clear. However, since we have ruled out changes in telomerase transcription and activity, it may act through blocking telomerase access to the telomere, by modification of a telomere binding protein or regulation of telomere protein levels.

### BRD4 inhibition shortens telomeres in long-term growth assays

The short-term lentiviral screen is a powerful, rapid method to identify inhibitors that block telomere elongation. To examine whether inhibition of BRD4 causes telomere shortening in a more physiological setting, we cultured cells for 6 weeks and examined telomere length by Southern blot. We chose OTX015 to inhibit BRD4 because it showed the greatest inhibition in blocking telomere elongation. OTX015 is also of particular interest because it is currently being tested in clinical trials to treat acute leukemia (57), lymphoma and multiple myeloma (58). Treatment with OTX015 caused significant telomere shortening in both human HeLa cells (Figure 4A), and mouse CAST/EiJ fibroblasts (Figure 4B). DMSO treated cells showed no change in telomere length over the same period. Notably, significant changes in the average telomere length were seen at 4 weeks in human cells, and after 2 weeks in mouse cells. The rapid nature of this telomere shortening suggests a relatively robust role for BRD4 in telomere maintenance.

**Figure 4.**
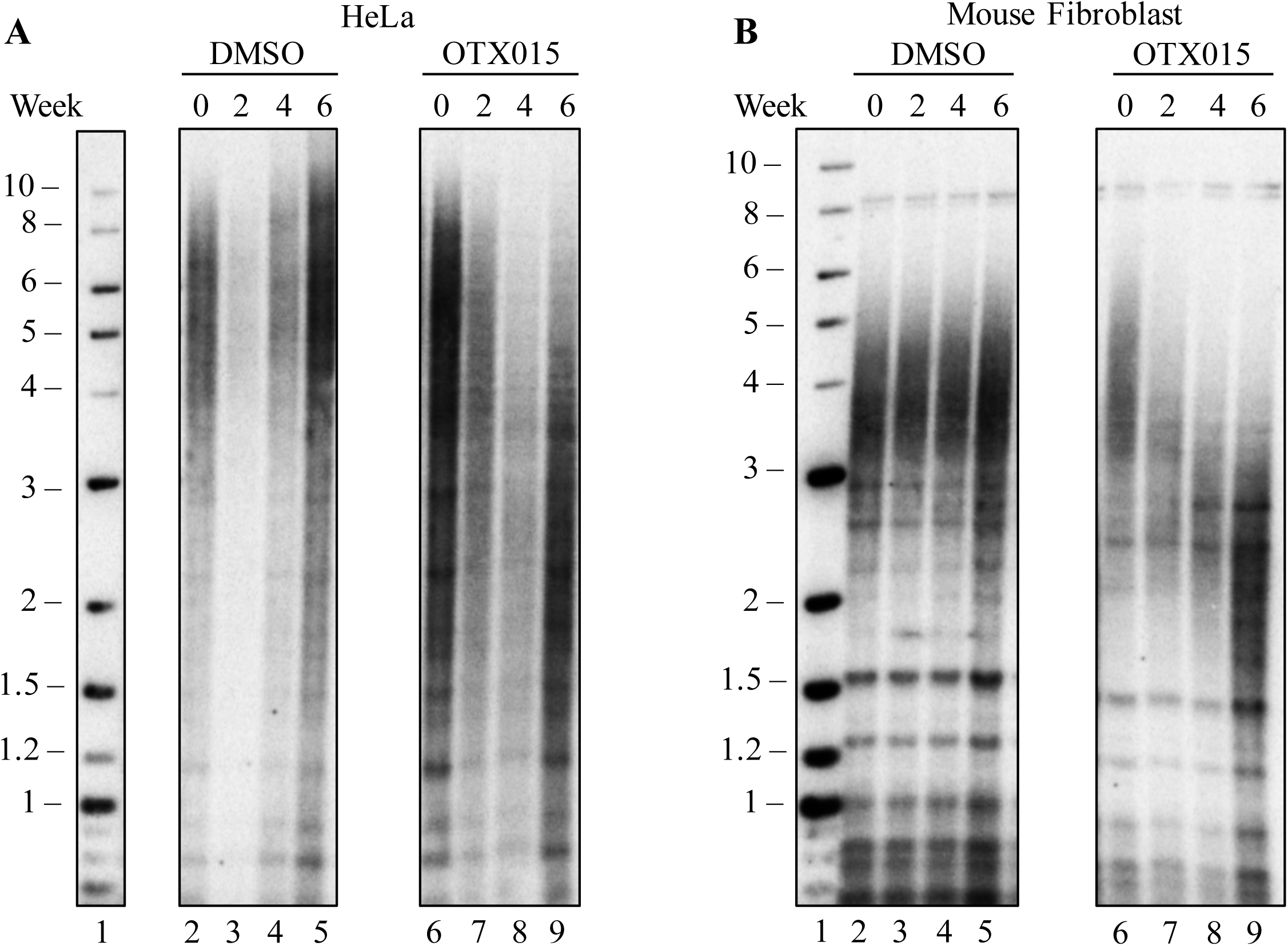
BRD4 inhibition causes telomere shortening in human and mouse cells in culture. (A) Southern blot of telomeric DNA from HeLa cells treated with DMSO (lane 2-5), or 2.5 μ M OTX015 (lane 6-9) for 6 weeks, with samples taken at 2, 4 and 6 weeks of treatment. (B) Southern blot of telomeric DNA from mouse fibroblast cells, which were treated with DMSO (lane 2-5), or 0.5 μ M OTX015 (lane 6-9) for 6 weeks, with samples taken at 2, 4 and 6 weeks of treatment.

BRD4 translocations are linked to cancer, and BRD4 inhibition reduces cell proliferation in acute myeloid leukemia (59,60), potentially by blocking BRD4 mediated transcription of c-MYC. Our finding, that BRD4 inhibition shortens telomeres in human and mouse cells in culture, suggests that part of the mechanism by which BRD4 inhibitors block cancer cell growth (60-62) may be through telomere shortening.

BRD4 inhibitors are currently in early phase clinical trials for treatment of hematopoietic cancers, including AML, ALL, multiple myeloma, and lymphoma (63).If telomere shortening also occurs *in vivo* in a clinical trial setting, telomere shortening might accentuate the anti-cancer effects of BRD4. BRD4 also affects immune cell function through its interaction with NF-**K**B (64), and BRD4 inhibitors have been proposed as anti-inflammatory agents, including in liver fibrosis and idiopathic pulmonary fibrosis (65,66). However, these approaches should be carefully considered, because short telomeres in humans lead to Telomere Syndromes that can manifest as bone marrow failure, pulmonary fibrosis, liver fibrosis, and other disease (67-69). Telomere shortening in either cancer trials or fibrosis may exacerbate underlying short telomere syndromes. Stratifying patients by telomere length before treatment may help identify individuals who might be at risk from side effects caused by further telomere shortening.

## ACKNOWLEDGEMENTS

We would like to thank Hao Zhang at the Johns Hopkins School of Public Health Cell Sorting Facility, and John Weger at the UC Riverside High Throughput Sequencing Center for many helpful discussions on the design and application of FACS and Illumina sequencing.

---

**Supplemental Table 1.** Compounds tested for effect on telomere elongation. The complete set of 36 compounds tested for their ability to block telomere elongation, when telomerase is overexpressed, is listed.

| **Number** | **Compound** | **Target** | **Vendor** | **Catalogue Number** |
| --- | --- | --- | --- | --- |
| 1 | KU-55933 | ATM | R&D Systems | 3544 |
| 2 | JQ1 | BRD4 | Selleckchem | S7110 |
| 3 | IBET151 | BRD4 | Selleckchem | S2780 |
| 4 | MS436 | BRD4 | Selleckchem | S7305 |
| 5 | OTX015 | BRD4 | Selleckchem | S7360 |
| 6 | MK-2206 2HCl | AKT | Selleckchem | S1078 |
| 7 | BI 2536 | PLK1 | Selleckchem | S1109 |
| 8 | Sorafenib | VEGFR, RAF | Selleckchem | S1040 |
| 9 | Staurosporine | PKC | Selleckchem | S1421 |
| 10 | Ruxolitinib | JAK1/2 | Selleckchem | S1378 |
| 11 | Rapamycin | mTOR | Selleckchem | S1039 |
| 12 | Selumetinib | ERK/MEK | Selleckchem | S1008 |
| 13 | CHIR-99021 | GSK-3$\alpha/\beta$ | Selleckchem | S2924 |
| 14 | Cerdulatinib | TYK2 | Selleckchem | S7634 |
| 15 | TG003 | CLK | Selleckchem | S7320 |
| 16 | WZ4003 | NUAK | Selleckchem | S7317 |
| 17 | SCH772984 | ERK1/2 | Selleckchem | S7101 |
| 18 | PD98059 | MEK | Selleckchem | S1177 |
| 19 | Trametinib | MEK1/2 | Selleckchem | S2673 |
| 20 | ML141 | CDC42 | Selleckchem | S7686 |
| 21 | Enzastaurin | PKC | Selleckchem | S1055 |
| 22 | BI-D1870 | RSK1/2/3/4 | Selleckchem | S2843 |
| 23 | KN-93 Phosphate | CaMKII | Selleckchem | S7423 |
| 24 | D 4476 | CK1 | Selleckchem | S7642 |
| 25 | CID755673 | PKD1/2/3 | Selleckchem | S7188 |
| 26 | 5-Iodotubercidin | CK1 | Selleckchem | S8314 |
| 27 | IC261 | CK1 | Selleckchem | S8237 |
| 28 | GDC-0994 | ERK1/2 | Selleckchem | S7554 |
| 29 | FR 180204 | ERK | Selleckchem | S7524 |
| 30 | Alisertib | AURKA | Selleckchem | S1133 |
| 31 | LDC000067 | CDK9 | Selleckchem | S7461 |
| 32 | SNS-032 | CDK2/7/9 | Selleckchem | S1145 |
| 33 | H 89 2HCl | PKA | Selleckchem | S1582 |
| 34 | SB203580 | p38 MAPK | Selleckchem | S1076 |
| 35 | BAY 80-6946 | PI3K | Selleckchem | S2802 |
| 36 | HTH-01-015 | NUAK1 | Selleckchem | S7318 |

